# ADELLE: A global testing method for Trans-eQTL mapping

**DOI:** 10.1101/2024.02.24.581871

**Authors:** Takintayo Akinbiyi, Mary Sara McPeek, Mark Abney

**Author notes:** Deceased (Takintayo Akinbiyi).

## Abstract

Understanding the genetic regulatory mechanisms of gene expression is a challenging and ongoing problem. Genetic variants that are associated with expression levels are readily identified when they are proximal to the gene (i.e., cis-eQTLs), but SNPs distant from the gene whose expression levels they are associated with (i.e., trans-eQTLs) have been much more difficult to discover, even though they account for a majority of the heritability in gene expression levels. A major impediment to the identification of more trans-eQTLs is the lack of statistical methods that are powerful enough to overcome the obstacles of small effect sizes and large multiple testing burden of trans-eQTL mapping. Here, we propose ADELLE, a powerful statistical testing framework that requires only summary statistics and is designed to be most sensitive to SNPs that are associated with multiple gene expression levels, a characteristic of many trans-eQTLs. In simulations, we show that for detecting SNPs that are associated with 0.1%–2% of 10,000 traits, among the 7 methods we consider ADELLE is clearly the most powerful overall, with either the highest power or power not significantly different from the highest for all settings in that range. We apply ADELLE to a mouse advanced intercross line data set and show its ability to find trans-eQTLs that were not significant under a standard analysis. This demonstrates that ADELLE is a powerful tool at uncovering trans regulators of genetic expression.

## Introduction

eQTL mapping, in which association is tested between gene expression levels and genetic variants, is a useful approach toward understanding mechanisms of genetic regulation. Cis-eQTLs, genetic variants that influence expression of proximal genes, are often readily detected because their effect sizes are commonly large, and the local nature of their effects limits the number of tests and, hence, the multiple testing burden. Because of this, many studies have focused on investigating the role of cis-regulatory effects on gene expression. Recent work, however, has estimated that cis-genetic effects account for a minority of human complex trait variance, perhaps as little as 11%, while trans-genetic effects, i.e. causes that are distant from the gene being regulated, may account for 70% or more of complex trait variance in humans [1, 2]. Unfortunately, even though trans-eQTL effects may dominate the genetic variability of gene expression and of complex traits, the identification of trans-eQTLs has been impeded by two significant hurdles. Compared to cis-eQTLs, trans-eQTLs are much harder to detect because their effect sizes tend to be smaller [2], and the space of possible genes whose expression they might be associated with is much bigger, leading to a higher burden of multiple comparisons.

A basic approach in both model organisms and humans to detect trans-eQTLs is to perform, for each SNP, a test of association against every trans-gene [1, 3–5]. To account for multiple testing, either a Bonferroni correction is applied or a false discovery rate (FDR) procedure is used. Because of the very high number of tests performed, only the strongest of signals achieve statistical significance. This has led to recent efforts to develop methods that will be more effective at detecting trans-eQTLs. Broadly, many of the methods seek to increase the number of discoveries by applying at least one of the following strategies (1) reducing the multiple testing burden by either reducing the number of variants tested [6–10] or reducing the number of genes tested [11–13], or (2) leveraging the expectation that a trans-eQTL will influence the regulation of multiple genes [12–15]. Although incorporating biological or other external information to effectively make the number of tests smaller has the potential to increase power by eliminating either variants or traits where the null hypothesis is true, it also has the potential to miss important signals. On the other hand, even though a trans-eQTL may affect the expression levels of multiple genes the number of these genes will typically be a very small fraction of the total number of genes. Together, these qualities have made the development of effective tools for the discovery of trans-eQTLs very challenging.

We address the problem of developing a powerful statistical method for trans-eQTL detection. In particular, we frame the problem as one where we seek to reject the global null hypothesis that for a candidate trans-eQTL (e.g., a single SNP) none of the expression traits are associated with the SNP. We develop a method that requires only summary statistics of individual tests of association between a SNP and an expression trait. Advantages of only requiring summary statistics include their ease of being shared and savings in the person and computational effort to generate them.

For the general statistical problem of aggregating a collection of *Z* scores or *p*-values into a single test of the global null hypothesis, various methods have been proposed. Examples include Simes’s method [16], Cauchy p-value combination [17], higher criticism [18, 19], the Berk-Jones statistic [20], and methods based on equal local levels (ELL) [20–25]. Both the higher criticism and Berk-Jones statistics have generalizations to the case where the tests are dependent, generalized higher criticism [26] and generalized Berk-Jones [27]. These methods were used to test association between a SNP-set and an outcome. Another class of global tests commonly used in genetics corresponds to the sum of *χ*^2^ statistics from different tests [28], which we call Sum-*χ*^2^. Variations and generalizations of this approach underlie methods for rare variant and haplotype association analysis such as SKAT [29] and other variance component tests [30]. The CPMA [14, 31] method has been proposed for combining test statistics for multi-trait mapping. The most commonly-used p-value combination approach is what we call Min-*p*, which is simply based on the smallest p-value in the collection, with significance assessed by Bonferroni correction or another approach such as Monte Carlo.

In general, there is no uniformly most powerful test of the global null hypothesis. Instead, different tests will be optimal in different alternative model regimes. For instance, the Min-*p* test, with a multiple testing correction, should do well when there is at least one extremely strong signal among the *p*-values. On the other hand, sums of χ^2^ types of tests are likely to do well when weak signals are spread over a relatively large proportion of the *p*-values [18, 26]. Here, we propose ADELLE, which is an extension of ELL to the case of dependent tests. Because ADELLE is an ELL-based test, we expect it to show strong performance when the signal is both relatively weak and sparse within a collection of *p*-values, which is the situation we expect when searching for trans-eQTLs. We assess the performance of ADELLE relative to other methods through simulation studies and application to trans-eQTL detection in mouse data from an advanced intercross line [4].

### Description of the method

We first briefly consider the simplified case in which the expression traits are assumed to be independent and describe how the ELL global testing method could be applied. Then we describe ADELLE, our extension of ELL to the case of dependent traits, which we apply to trans-eQTL mapping.

### Global trans-eQTL testing with ELL

In an eQTL mapping study in which 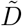 expression traits and *M* genome-wide SNPs are observed on each of *N* individuals, suppose each expression trait is tested for association with each genome-wide SNP in the sample leading to a 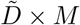 summary statistic matrix Π of p-values having (*d, m*)th entry *π*_*dm*_ equal to the p-value for testing association between expression trait *d* and SNP *m* in the *N* individuals. In this subsection we make the simplifying assumption that the 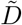 traits are independent. We extend to the case of dependent traits in the following subsection.

For a given SNP *m*, define 𝒟_*m*_ to be the subset of expression traits that are considered trans to it, from among the larger set of 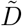 traits measured. To detect trans-eQTLs, we propose to perform *M* global hypothesis tests, one for each SNP, in which the *m*th global hypothesis test has null and alternative hypotheses

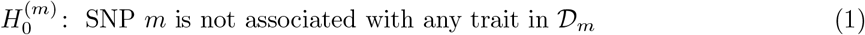

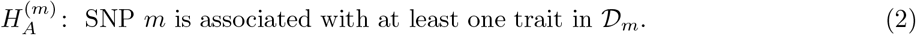

We now fix a SNP *m* and describe the ELL method for performing the *m*th global hypothesis test, where the test statistic is constructed from the p-values in column *m* of Π. Specifically, we consider a vector of p-values *π* of length *D*_*m*_ = |𝒟_*m*_|, consisting of the subset of p-values in the *m*th column of Π that correspond to the traits in 𝒟_*m*_. For simplicity of exposition, we drop the subscript *m* in the remainder of this subsection, so we consider 𝒟 to be the set of traits that are trans to the SNP and consider *π* to be of length *D*. Under the null hypothesis that the given SNP is not associated with any of its trans traits and the further assumption of independence of traits (and assuming that the method for calculating p-values is well-calibrated), the entries of *π* would be *D* independent and identically distributed (i.i.d.) Uniform(0,1) random variables.

ELL is a general global testing method that models the entries of *π* as i.i.d. from a distribution having cumulative distribution function (cdf) *F*_*π*_(*x*) for *x* ∈ (0, 1). The null hypothesis would be

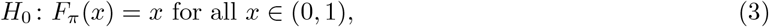

i.e., the p-values are Uniform(0,1), and the one-sided alternative hypothesis would be

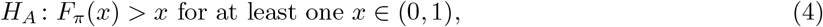

i.e., the p-values tend to be smaller under the alternative than would expected under the null. We use the notation *π* = (*π*_1_, …, *π*_*D*_) and for 1 ≤ *d* ≤ *D*, we define *π*_(*d*)_ to be the *d*th order statistic of *π*, i.e., we sort the entries of *π* in ascending order and let *π*_(*d*)_ be the *d*th component of the sorted vector, so *π*_(1)_ ≤ *π*_(2)_ ≤ … ≤ *π*_(*D*)_. Under the null hypothesis that the unsorted p-values *π*_1_, …, *π*_*D*_ are i.i.d. uniform, the entries of (*π*_(1)_, …, *π*_(*D*)_) are dependent with a known joint distribution, and marginally each *π*_(*d*)_ has the Beta(*d, D* − *d* + 1) distribution for 1 ≤ *d* ≤ *D*.

The ELL test starts by comparing each order statistic to its corresponding beta null distribution and deciding whether it is smaller than expected. Then the ELL test statistic is based on the order statistic that shows the most significant deviation from its corresponding null distribution. On the one hand, if trans-eQTL signals are only of moderate or weak size, then, e.g., *π*_(1)_ and *π*_(2)_ might actually represent null tests, and the true alternatives could be represented by smaller than expected *π*_(*d*)_ for values of *d* that are perhaps of small to moderate size. On the other hand, finding that *π*_(*d*)_ is smaller than expected only for larger values of *d*, e.g., *d* close to *D*, would be difficult to interpret and might not seem compelling evidence for the SNP being a trans-eQTL. Therefore, we propose to base the ELL test statistic on only the smallest fraction *q* of the p-values, i.e., on order statistics *π*_(*d*)_ for 1 ≤ *d* ≤ *qD*, where *q* ∈ (0, 1). In the original formulation of ELL, Berk and Jones [20] used *q* = .5. In the eQTL mapping context, a smaller *q* would seem more appropriate, and we take *q* = .2, i.e., we only the consider the smallest 20% of the p-values for a given SNP. For simplicity of notation, in what follows we assume that *qD* turns out to be an integer (otherwise it could be replaced by floor(*qD*)).

To construct the ELL test statistic, we first calculate *qD* “l-values”, one for each *π*_(*d*)_, 1 ≤ *d* ≤ *qD*, where the l-value *l*_*d*_ for *π*_(*d*)_ is the p-value for testing the null hypothesis that *π*_(*d*)_ is drawn from a Beta(*d, D* − *d* + 1) distribution vs. a one-sided alternative for which we reject the null hypothesis if *π*_(*d*)_ is sufficiently small. Thus, *l*_*d*_ = pbeta(*π*_(*d*)_, *d, D* − *d* + 1) where pbeta(*x, a, b*) is the cdf of the Beta(*a, b*) distribution evaluated at *x*. Then the ELL test statistic is

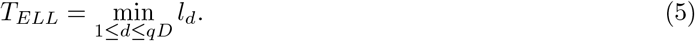

To assess whether the SNP is a trans-eQTL, we perform a one-sided hypothesis test at level *α* based on *T*_*ELL*_, where we reject the null hypothesis in Eq 1 if *T*_*ELL*_ *< η*, where *η* (the “local level”) is a function of *α*. We refer to this as an equal local level test because the local level *η* at which we reject *H*_0_ is equal for all *l*_*d*_. That is, if any of the l-values are less than *η* we reject *H*_0_. Previous work [24] shows that the ELL test is asymptotically optimal for detecting deviations from a Gaussian distribution for a wide class of rare-weak contamination models.

For the case when the traits are independent, there are existing algorithms [25, 32, 33] to calculate the global level *α* of the test as a function of the local level *η*, where we call this function *α*(*η*). These algorithms are specifically for the case *q* = 1. However, we have adapted the algorithm of Weine et al. 2003 [25] to more general *q*. To do this, we let *ξ* = floor(*qD*), and then obtain *α*(*η*) as 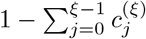, where 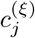 is a quantity calculated recursively in Algorithm 1 of Appendix B.2 of Weine et al. [25] To invert the function *α*(*η*) and determine the local level *η* corresponding to a chosen global level *α* for the ELL test, we conduct a binary search to find the needed *η*.

### ADELLE: extension of the ELL method to dependent traits

The ELL approach described in the previous subsection assumes independence of traits, but in practice there is typically correlation among gene transcript levels. Our goal is still to perform, for each SNP, a global test based on the null and alternative hypotheses in Eq 3 and 4. However, dependence among traits leads to dependence among the elements of the p-value vector *π*. In that case, it is no longer true that, e.g., *π*_(*d*)_ is beta distributed under the null as it is in the independence case. Therefore, the methods we describe above for calculation of the ELL test statistic and its null distribution are no longer applicable.

The ADELLE method we propose generalizes the ELL approach to allow for dependent traits. For 1 ≤ *d* ≤ *D*, define *F*_(*d*)_ to be the cdf of the distribution for *π*_(*d*)_ under the null hypothesis in the case when the traits are dependent. The basic idea behind ADELLE is that we find an approximation to *F*_(*d*)_ and use it to calculate the *qD* l-values *l*_1_, …, *l*_*qD*_ in the case when the traits are dependent. Then we define the ADELLE test statistic *T*_*ADELLE*_ to be the minimum of *{l*_1_, …, *l*_*qD*_*}*. Finally, we calculate the p-value for the ADELLE test using a Monte Carlo approximation method given in subsection **Monte Carlo p-value calculation**.

First we describe how dependence is incorporated into the model. Rather than directly modeling the dependence on the p-value scale, we instead consider a set of association test statistics *Z*_1_, … *Z*_*D*_, where *Z*_*d*_ tests association between the given SNP and its *d*th trans trait, 1 ≤ *d* ≤ *D*. We assume that under the null hypothesis, each *Z*_*d*_ ∼ *N* (0, 1), where they can be correlated with each other, and we assume that *π*_*d*_ is a two-sided p-value based on *Z*_*d*_, i.e., *π*_*d*_ = 2Φ(−|*Z*_*d*_|), where Φ is the standard normal cdf.

Let *G* denote the genotype vector of the SNP and *Y*_*d*_ the phenotype vector of its *d*th trans trait. Typical examples of *Z*_*d*_ would be the t-statistic for testing significance of *G* in a linear model for *Y*_*d*_ or the Wald t-statistic for testing significance of *G* in a linear mixed model (LMM) for *Y*_*d*_. In large samples, such a t-statistic will be approximately standard normal under the null hypothesis or, if necessary, could be transformed to be approximately standard normal under the null hypothesis by applying the transformation Φ^−1^(pt(*Z*_*d*_)) where pt is the cdf of the t-distribution with degrees of freedom = *N* − *k* − 1 where *k* is the number of predictors in addition to the intercept in the linear model or LMM. A likelihood ratio *χ*^2^ test statistic for testing significance of *G* in a LMM for *Y*_*d*_ could also be converted to such a *Z*_*d*_ value by taking a square root of the 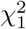 test statistic and applying the sign of the estimated coefficient of *G* in the LMM for *Y*_*d*_.

We let *Z* = (*Z*_1_, …, *Z*_*D*_)^*T*^ and, under the global null hypothesis that the SNP is unassociated with any of its trans traits, we model *Z* as multivariate normal:

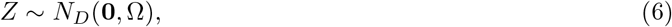

where *N*_*D*_ denotes the multivariate normal distribution of dimension *D*, **0** is a vector of 0’s of length *D* and Ω is the *D × D* trait correlation matrix. (See Supplementary Information for a derivation of this model.) To estimate Ω, we first form the sample correlation matrix *Ĉ*_*trait*_ for the 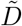 traits. However, the estimate *Ĉ*_*trait*_ would be low rank because typically 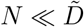, so we could regularize it by using the shrinkage estimator [34] 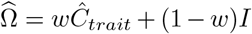. (See Supplementary Information for details on choice of the regularization parameter *w*.)

To calculate *F*_(*d*)_(*h*) for *h* ∈ (0, 1), where *F*_(*d*)_ is the cdf of *π*_(*d*)_ under the null hypothesis, we first point out the key identity that the two events *E*_1_ = *{π*_(*d*)_ ≤ *h}* and 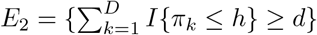 are the same, where *I{·}* is the indicator function that equals 1 if the event inside the brackets occurs and 0 otherwise, and where *E*_2_ is saying that at least *d* of the p-values are ≤ *h*. By the defined relationship between *π*_*k*_ and *Z*_*k*_, we have that the events *{π*_*k*_ ≤ *h}* and *{*|*Z*_*k*_| ≥ −Φ^−1^(*h/*2)*}* are the same, so 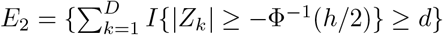. Next, define 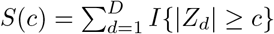 for *c* ≥ 0, where *S*(*c*) counts the number of |*Z*_*d*_| that are greater than or equal to *c*, and note that *E*_2_ = *{S*(−Φ^−1^(*h/*2)) ≥ *d}*. Therefore, the following two events are the same

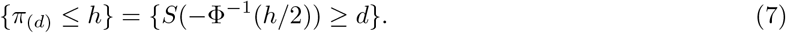

Finally, we have for the *l*-value

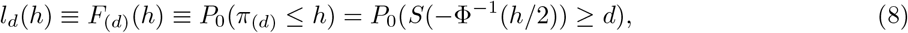

where *P*_0_(*·*) represents probability under the null hypothesis that the SNP is not associated with any of its *D* trans traits.

As a consequence, we can obtain needed values of *F*_(*d*)_ by considering the distribution of *S*(*c*) under the null hypothesis. If Ω = *I*, then for *c* ≥ 0, *S*(*c*) has the null distribution of a Binomial(*D*, 2Φ(−*c*)) random variable. When Ω *≠ I, S*(*c*) has the same null mean as a Binomial(*D*, 2Φ(−*c*)), but the null variance of *S*(*c*) is strictly greater than that for Binomial(*D*, 2Φ(−*c*)), i.e., the distribution of *S*(*c*) is over-dispersed relative to binomial. The beta-binomial distribution is a standard choice for modeling binomial-like data when there is over-dispersion. Therefore we approximate the distribution of *S*(*c*) with a beta-binomial distribution *BB*(*D, λ, γ*) where *λ* and *γ* are chosen so that the first and second moments match those of *S*(*c*), using techniques of a previous work [35] (see also [36]). The details are given in Supplementary Information. From the resulting approximation to the distribution of *S*(*c*), we obtain an approximation to *F*_(*d*)_, which we call 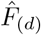, based on Eq. 8. The required calculation of 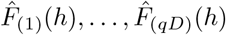 for all *h* ∈ *ℋ* can be efficiently carried out as a pre-computation step, as described in detail in Supplementary Information.

To obtain the ADELLE test statistic, we first obtain the *qD* l-values *l*_1_, …, *l*_*qD*_, where *l*_*d*_ is defined to be 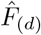 evaluated at the observed value of *π*_(*d*)_. Then the ADELLE test statistic is given by *T*_*ADELLE*_ = min_1≤*d*≤*qD*_ *l*_*d*_. In the special case when Ω = *I*, we get back the same ELL l-values and ELL test statistic used for the independence case in the previous subsection.

### Monte Carlo p-value calculation

We use a Monte Carlo approach to assess significance of the ADELLE global test statistic. Specifically, we simulate *R* i.i.d. vectors 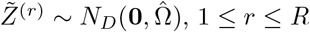, 1 ≤ *r* ≤ *R*, where *R* is very large (e.g., 10^7^ in the data analysis), and for each 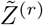, we calculate the ADELLE statistic, call it *T* ^(*r*)^. For any observed ADELLE statistic, *T*, we calculate its p-value as (*N* (*T*) + 1)*/*(*R* + 1), where 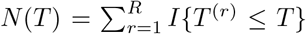 counts the number of *T* ^(*r*)^ values that are less than or equal to *T*. We use the same Monte Carlo method to assess significance of the G-Null, CPMA and Sum-χ^2^ global test statistics in our simulation studies, where these three global test statistics are described below in subsection **Additional global test statistics included in the comparison**. In the simulations, we verify the empirical type 1 error of our Monte Carlo p-value calculation for the ADELLE, G-Null, CPMA and Sum-χ^2^ global testing methods.

### Simulation methods

In the simulations, we consider a setting in which we have summary statistics from association tests of a SNP with each of *D* = 10^4^ expression traits, and we want to combine the summary statistics into a global test of the null hypothesis that the SNP is not associated with any of the traits. We use the ADELLE method and each of the 6 different global testing methods described below in subsection **Additional global test statistics included in the comparison** to perform the global test. To assess type 1 error at level *α*, we generate *r* = 10^4^ simulation replicates in which the SNP is not associated with any of the *D* traits and calculate each of the global test statistics on each replicate. For each *α* level and each global testing method, we estimate type 1 error by the proportion of replicates in which the given global testing method produced a p-value *< α*. To compare power across methods, we generate 10^3^ simulation replicates in which the SNP is associated with exactly *A* of the *D* traits, where we perform studies for each of several choices of *A* from 5 to 200. The effect size of the SNP on each of the *A* associated traits is set to be *c*_*A*_, where *c*_*A*_ is chosen so that the maximum power across the methods is approximately in the range 0.5–0.9. We compare the power of the different methods based on the proportion of replicates in which each method rejects the null hypothesis. To simulate the data, we start by randomly choosing a true trait correlation matrix Ω (see Supplementary Information for details). To perform the Monte Carlo p-value calculation described in the previous subsection, we simulate trait values for the 10^4^ traits for 10^3^ individuals, from which we estimate 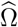 as described in subsection **ADELLE: extension of the ELL method to dependent traits** above. The estimated 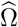 is then used in the Monte Carlo p-value calculation. In each simulation replicate, we simulate a vector of *Z* scores of length *D* from a multivariate normal distribution with mean vector *µ* = 0 under the null hypothesis and with correlation matrix Ω. Under the alternative hypothesis, we simulate the *Z* scores from the same distribution as under the null hypothesis but where the mean vector has exactly *A* of the *D* entries equal to *c*_*A*_ and the remaining *D* − *A* entries equal to 0.

### Additional global test statistics included in the comparison

We assessed the type 1 error and power of ADELLE as well as the following methods for testing the global null hypothesis that a given SNP is not associated with any expression trait. For each replicate a vector of (dependent) *Z* scores was generated as described above and given as input to each method.

**Min-***p* For each *Z* score vector, to obtain its p-value using the Min-*p* method, we calculate *π*_(1)_*D*, the Bonferroni-corrected minimum p-value among all the test statistics in the vector *Z*.

**Simes** For each *Z* score vector, to obtain its p-value using the Simes method [16], we calculate min 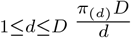. The Simes p-value is closely related to the Benjamini-Hochberg procedure [37] for controlling FDR.

**Cauchy** For each *Z* score vector, to obtain its p-value using the Cauchy method [17], we calculate 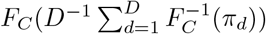, where *F*_*C*_ is the Cauchy cdf.

**G-Null** The G-Null method is a simpler variation on the ADELLE method. In the ADELLE method, the estimated trait correlation matrix 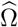 is used both in (1) calculating the *l*-values used to construct the test statistic and in (2) the Monte Carlo p-value calculation. In contrast, in the G-Null method, the *l*-values are calculated assuming independence, and 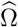 is used only for the Monte Carlo p-value calculation. As a result, both methods would be expected to have correct type 1 error, and ADELLE would be expected to have higher power when there is dependence among the traits. In simulations, we investigate to what extent using 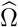 to calculate the *l*-values allows ADELLE to improve power over G-Null.

For each *Z* score vector, to obtain its p-value using the G-Null method, we first calculate the ELL test statistic given in Eq 5. If the elements of *Z* were independent, we could calculate a p-value by the method given in subsection **Global trans-eQTL testing with ELL**. However, because they are dependent, we instead obtain a Monte Carlo p-value using the method described in subsection **Monte Carlo p-value calculation** above.

**Sum-χ**^2^ The test statistic is 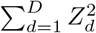 the sum of the squares of the *Z* scores in the vector. If the *Z* scores were independent under the null hypothesis, this test statistic would be 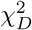 distributed. However, in this setting they are dependent, and we instead obtain a Monte Carlo p-value using the method described in subsection **Monte Carlo p-value calculation** above.

**CPMA** We used our own implementation of the method described in [14] to compute the CPMA statistic. The CPMA test models the elements of the vector (− log(*π*_1_), .., log(*π*_*D*_)) as i.i.d. draws from an Exponential(*λ*) distribution and tests the null hypothesis *λ* = 1 vs. the alternative *λ ≠* 1. We compute the likelihood ratio statistic for this test. Because the chi-squared null distribution does not hold when the *p*-values are correlated, we instead used the Monte Carlo p-value calculation described above.

In addition we considered both the GHC [26] and GBJ [27] methods but were unable to successfully run the available software on the scale of problems we consider here.

### Trans eQTLs in an advanced intercross line

Gonzales et al. [4] described an advanced intercross line (AIL) of mice and undertook genome-wide association studies (GWAS) and eQTL mapping studies in this population. They report finding thousands of cis and trans eQTLs across three brain regions. Here, we focus on trans eQTL associations in the hippocampus region and use summary statistics to test for trans eQTL associations that were not significant in the original study. Details of the data set and original analysis can be found in Gonzales et al. [4].

For expression traits in the hippocampus, Gonzales et al. determined that in their dataset a p-value threshold of 9.01 *×* 10^−6^ corresponded to genome-wide significance of 0.05 when correcting for SNP-wise multiple testing, based on a permutation analysis. This value of 9.01 *×* 10^−6^ would thus be an appropriate significance threshold for testing a single expression trait with all SNPs in the genome, and it would also be an appropriate threshold for a global testing method such as ADELLE or any of the other 6 methods described above, in which the p-values for a given SNP with each possible expression trait are combined into a single test statistic, resulting in one test performed for each SNP in the genome. However, if one instead takes a non-global-testing strategy of considering all the p-values for every possible pairing of a SNP and one of its trans traits, then in order to identify a SNP as a trans eQTL with a type 1 error rate of 0.05, it is necessary to correct for both the number of SNPs and the number of traits tested. For any SNP in this study there are approximately 14,000 trans genes against which it is tested. After doing a Bonferroni correction, we, therefore, consider a single SNP-trans gene association to be statistically significant if its p-value is less than 6.4 *×* 10^−10^.

ADELLE only requires summary statistics and a trait correlation matrix, but the available results for this data set only include summary statistics for associations that had *p*-value less than 9.01 *×* 10^−6^. To allow larger p-values to potentially contribute to the global test, we decided to regenerate the complete set of SNP-gene expression *Z* scores. We downloaded the G50–56 LGxSM AIL GWAS data set available at https://palmerlab.org, filtered the genotype dosage file to include only those mice that had gene expression data in the hippocampus, and pruned SNPs that were in complete LD using Plink [38], leaving 9671 SNPs across the genome. We used the downloaded gene expression matrix *Y* for the hippocampus that had all covariates regressed out and was quantile normalized. We computed the sample trait correlation matrix based on the 15,071 autosomal, expressed genes in *Y* and applied our regularization method to obtain 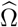. Following the code provided in the supplementary information of Gonzales et al. [4], we used the software package Gemma [39] to construct LOCO GRMs and to do association analysis between each SNP-gene expression pair, which is the equivalent of performing ∼ 15, 000 different GWASs. Using the Monte Carlo assessment of significance based on 10^7^ replicates, we determined an empiric ADELLE p-value for every SNP and used the genomewide significance cutoff of 9.01 *×* 10^−6^ that is needed to correct for SNP-wise multiple testing in this dataset.

## Results

### Power and type 1 error

We tested type 1 error at significance levels *α* = 0.05, 0.01, and 0.001. As seen in Table 1, all methods control the type 1 error rate at the nominal level. This validates the Monte Carlo p-value calculation approach that we use for ADELLE, G-Null, Sum-χ^2^ and CPMA.

**Table 1.** Type 1 error rates of different global testing methods.

| Method | Empirical type 1 error at nominal level |  |  |
| --- | --- | --- | --- |
|  | 0.05 | 0.01 | 0.001 |
| Adelle | 0.0498 | 0.0097 | 0.0006 |
| G-Null | 0.0488 | 0.0094 | 0.0007 |
| Min-P | 0.0487 | 0.0089 | 0.0004 |
| Simes | 0.0503 | 0.0089 | 0.0004 |
| Cauchy | 0.0512 | 0.0088 | 0.0004 |
| Sum-$\chi^2$ | 0.0485 | 0.0086 | 0.0007 |
| CPMA | 0.0466 | 0.0091 | 0.0007 |

Type 1 error is based on $10^4$ replicates in each case. The acceptance region for a test (at level 0.05) of whether the type 1 error rate differs from the nominal 0.01 level based on $10^4$ replicates is (0.00805, 0.01195). The corresponding acceptance regions for nominal levels 0.05 and 0.001 are (0.04573, .05427) and (.00038, .00162), respectively.

The results of the power simulations for all 7 methods considered are given in Tables S1 - S6 of Supplementary Information. Fig 1 shows the results for 6 of the 7 methods. (In order to make the plots less cluttered, the power of the Simes method was not included in Fig 1, as it was approximately equal to that of Min-P in all cases.) It is particularly illuminating to examine the relative power of the methods across different numbers of associated expression traits for the tested trans e-QTL, where this is shown in Fig 2. For each choice of the number of associated expression traits, we plot the power of each method divided by the maximum power observed across all the methods for that setting. We can see that the Min-P, Simes, and Cauchy methods behave all similarly. As expected, they perform best with a small handful of associated traits, e.g., in our simulations, these methods perform better than the other methods when 5 out of 10^4^ of the tested traits are associated. However, their power is significantly below that of ADELLE with 20 or more associated traits, and they are the worst performing methods with 100 or 200 associated traits. At the opposite end of the spectrum are the Sum-χ^2^ and CPMA methods, which perform the worst with a small handful of associated traits, but outperform Min-P, Simes and Cauchy with a very large number of associated traits. For the Sum-χ^2^ method, our results are consistent with previous observations about the performance of this type of method [17, 18, 26]. In contrast, the ADELLE method emerges as the most powerful method when there are a moderate number of associated traits. When the number of associated traits is in the range of 10 to 200, ADELLE’s power is either the highest or not significantly different from the highest, and it clearly dominates all the other methods in terms of power when the number of associated traits is in the range of 20 to 100. Overall, ADELLE is the only method that consistently maintained high relative power across the entire range of scenarios we tested (5 to 200 associated traits out of 10^4^).

**Fig 1.**
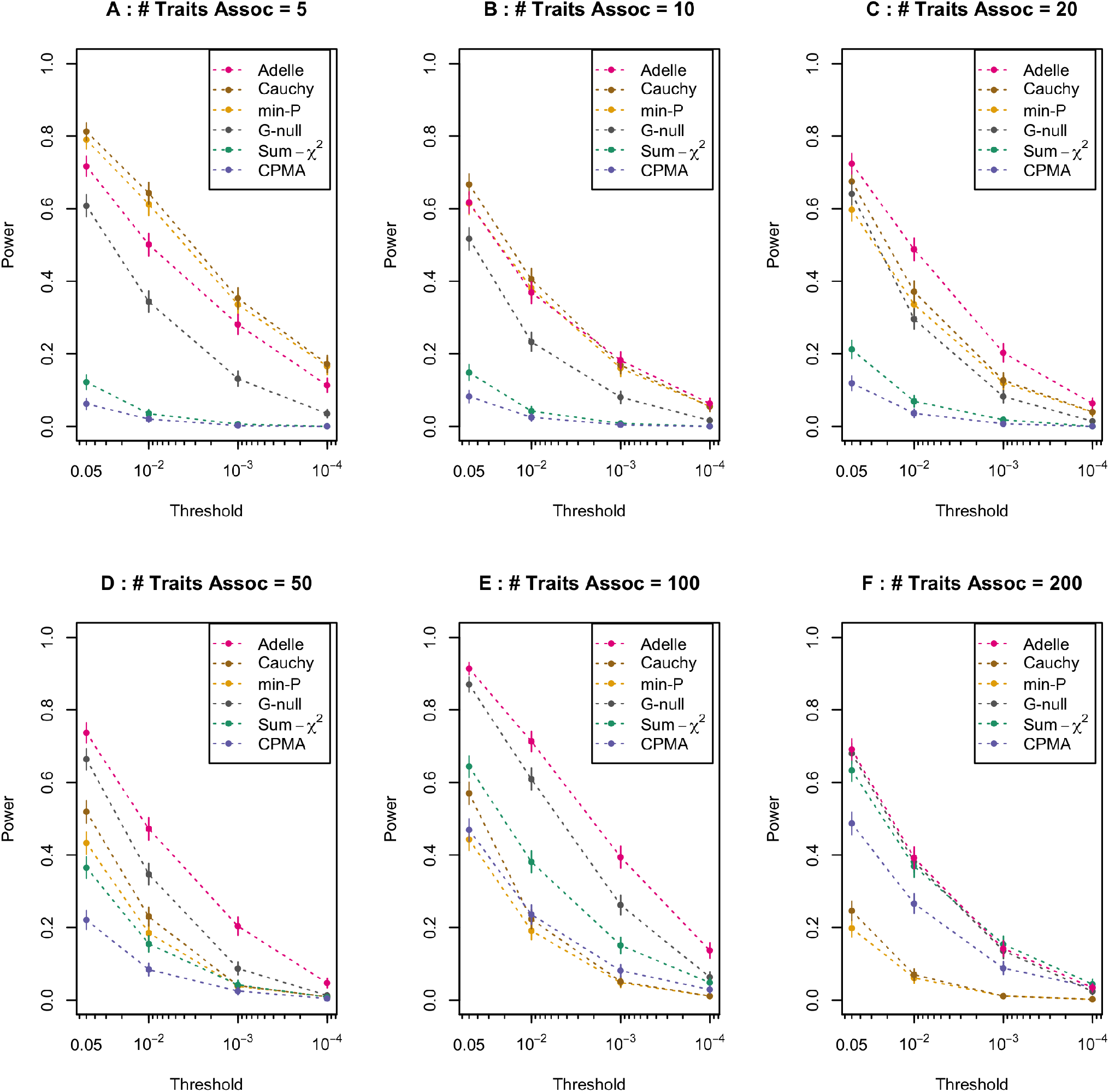
Power curves comparing different global testing methods for detecting a trans-eQTL. Each panel shows power for detecting a trans-eQTL plotted against the significance threshold of the association test, for 6 of the 7 global testing methods considered. In order to make the plots less cluttered, the power of the Simes method was not plotted, as it was approximately equal to that of Min-P in all cases (see Tables S1 - S6). In each panel, power is based on 10^3^ simulated replicates. Each panel shows the plot for a setting in which a given number of expression traits are associated with the tested trans-eQTL. For each point of the plot, the corresponding vertical bar represents the 95% confidence interval for power. In Panel A, the number of associated expression traits is 5. In Panels B, C, D, E, and F, the numbers of associated expression traits are 10, 20, 50, 100, and 200, respectively.

**Fig 2.**
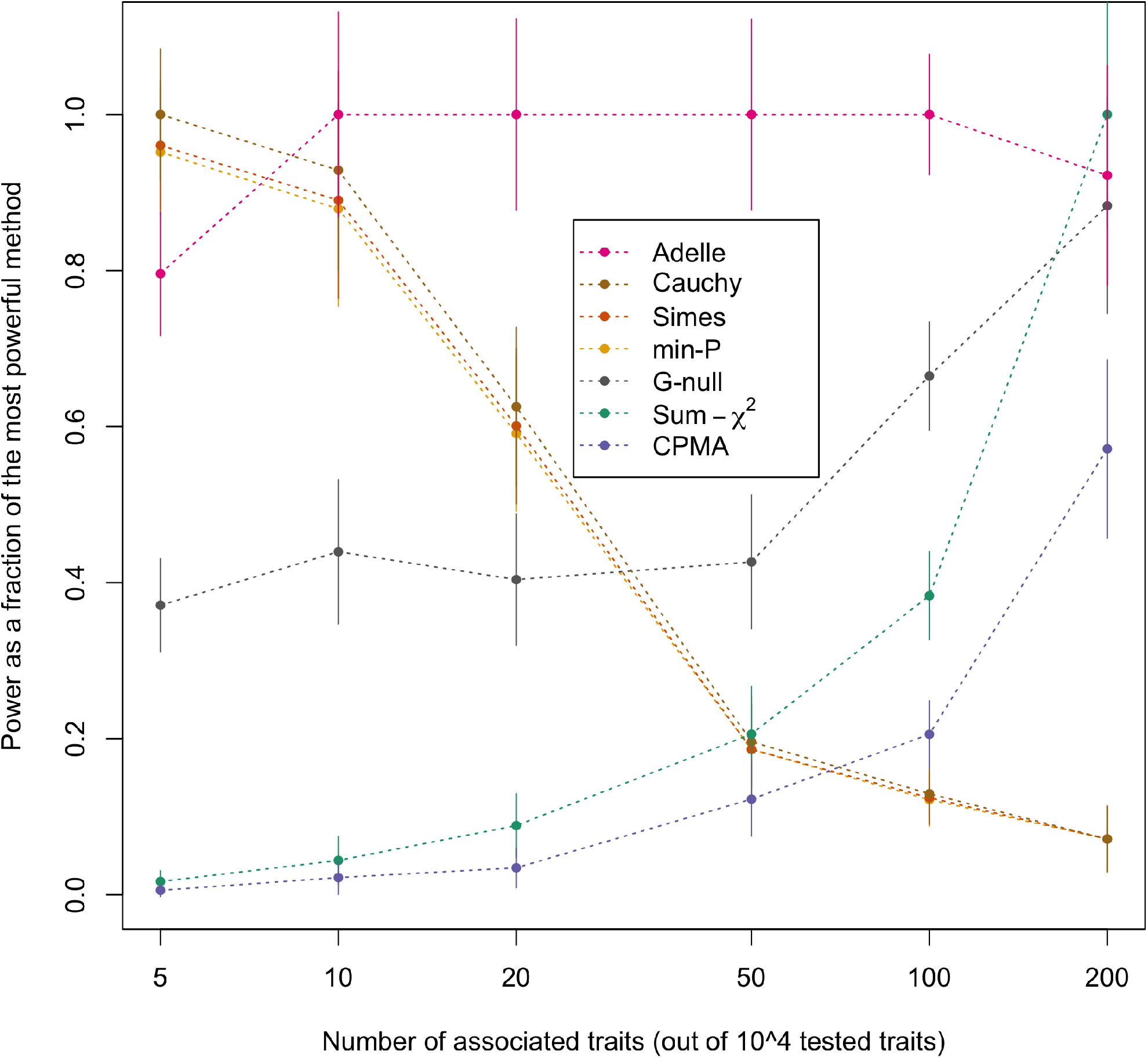
Relative power vs. number of associated traits for different global testing methods. Relative power at significance level 0.001, based on 10^3^ simulated replicates, is plotted against the number of associated traits out of 10^4^ total traits tested, for each of the 7 global testing methods considered. For a given number of associated traits, relative power for a method is defined as its power divided by the maximum power achieved by any of the 7 methods for that setting. For each point of the plot, the corresponding vertical bar represents the 95% confidence interval.

Inclusion of the G-Null method allows us to examine the effect of using the estimated trait correlation matrix 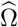 in calculating the *l*-values of the ADELLE statistic. The difference between G-Null and ADELLE is that G-Null uses an identity matrix in place of 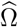 in calculating the *l*-values, which leads to a simple closed-form expression. However, from Figs 1 and 2, we can see that ADELLE has significantly greater power than G-Null for most scenarios. From this, we can conclude that it is important to use Ω in calculating the *l*-values.

### Trans eQTLs in an advanced intercross line

In the supplementary information to their article, Gonzales et al. [4] list all trans associations (where a “trans association” is defined to be any association signal that is detected between a SNP and an expression trait for a gene where the SNP and the gene are located on different chromosomes) in the hippocampus that had p-value less than 9.01 *×* 10^−6^, which corresponds to the threshold when correcting for all SNPs in the genome. Thus, many of the listed potential trans eQTLs do not meet the more stringent significance level of 6.4 *×* 10^−10^ required when correcting for both SNP-wise and trait-wise multiple testing.

Across the genome we replicated the trans eQTLs discovered by Gonzales et al. [4]. With the exception of one locus on chromosome 12, all trans eQTLs discovered with ADELLE also reached the significance threshold of 6.4 *×* 10^−10^ in the previous [4] analysis. However, in the region of chromosome 12 from 70 - 74 Mbp, shown in Fig 3, there are several SNPs that are detected as significant by ADELLE but are not detected as significant trans eQTLs by the previous analysis [4] when multiple testing is accounted for. In the previous analysis, these SNPs each show a sub-significant level of association across multiple expression traits. The five SNPs in this chromosome 12 region that were detected as significant by ADELLE are listed in Table S7 in the Supplementary Information.

**Fig 3.**
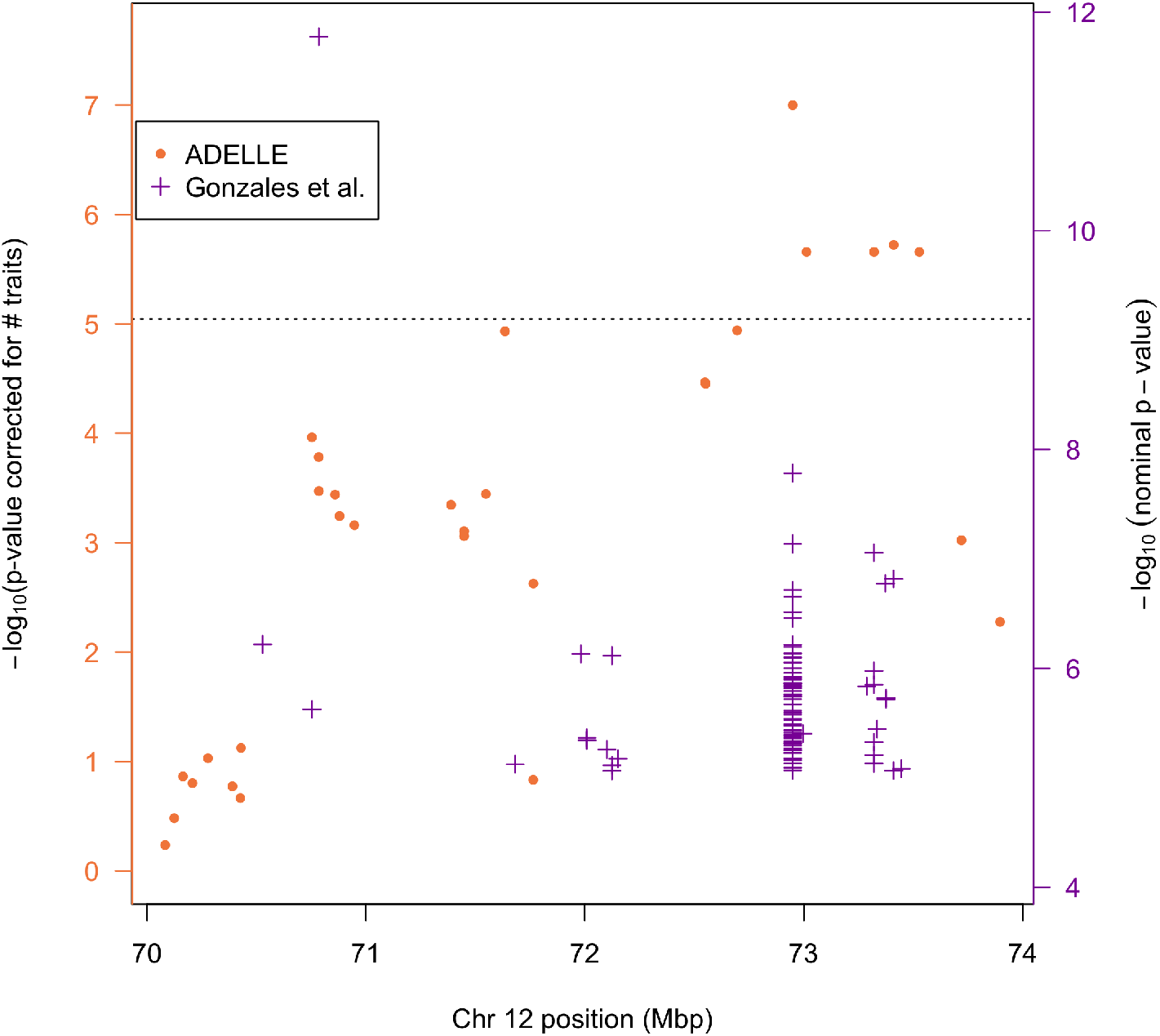
Trans eQTL associations in a region of chromosome 12. The purple “+” symbols in the figure represent single SNP-trait associations in the Gonzales et al. [4] analysis that had *p*-value less than 9.01 *×* 10^−6^. Among the purple crosses, a single SNP may appear multiple times with different *p*-values in the figure, representing tests of the same SNP with different traits. The − log_10_ of these *p*-values are displayed on the right-hand axis. The left-hand axis represents a Bonferroni-corrected version of the right-hand axis, in which a correction for testing 14,078 traits is made. The ADELLE global testing result for each SNP in the region is shown as an orange dot whose *p*-value on the − log_10_ scale is shown on the left-hand axis, because the ADELLE *p*-value already accounts for testing multiple traits. The dotted line represents the genomewide significance threshold, correcting for both multiple SNPs and multiple traits. Note that this dotted line is more stringent than the one used in Gonzales et al. because we have applied a Bonferroni correction for the number of traits (i.e. gene expressions) tested at each SNP.

In Fig 3, we can see both the ADELLE results and the previously reported results. Among the purple crosses (previous results), a single SNP may appear multiple times, with different *p*-values representing tests of the same SNP with different traits. The Min-*p* global testing result for a given SNP would be represented by the highest purple cross at a given location, with corresponding − log_10_ *p*-value given by the left-hand axis. The only SNP in this region that surpasses the threshold to be a trans eQTL in the previous analysis is at approximately 70.8 Mbp. This SNP is strongly associated with only a single trait, while its p-values for association with the remaining traits fit well to the uniform null distribution. In such a case, Min-*p* is expected to have high power. From Fig 3, we can see that the most significant result in the region by the ADELLE method (SNP rs262318378 at approximately 72.9 Mbp) had many small but sub-significant *p*-values for association in the previous analysis, and so is not significant by Min-*p*. This is a setting in which ADELLE is expected to have higher power. The other 4 significant ADELLE results in the region correspond to SNPs in high LD with rs262318378, so this set of 5 results may correspond to a single trans eQTL signal. Interestingly, for 2 of these 4 SNPs, the ADELLE test is significant, but but none of the individual trait *p*-values for these SNPs was small enough to be reported by Gonzales et al. (i.e., none pass the nominal 9.01e-6 threshhold). In other words, none of the individual SNP-trait *p*-values for these SNPs even meets the significance threshold when correcting only for SNP-wise multiple testing, much less the more stringent standard of correcting for both SNP-wise and trait-wise multiple testing. This occurs because of an enrichment of many small p-values at that SNP, but where none of these p-values by iself is smaller than 9.01e-6. There is also a SNP at approximately 71.6 Mbp (which is not in strong LD with the 5 SNPs having significant ADELLE p-values) that is nearly significant using ADELLE, but for which there is also no single expression trait whose p-value is even as small as 9.01e-6. The data analysis results are consistent with the simulation results that showed ADELLE can gain power for trans eQTL mapping in a setting in which there are a moderate number of associated traits with relatively weak effects.

This region on chromosome 12 with many small but not statistically significant p-values was previously noted [4] and referred to as a “master” eQTL. Trans eQTLs acting as master regulators, that is affecting the expression levels of many genes, have been observed previously [2, 40, 41] and may often be located in trans eQTL hot spots [41, 42]. One possible mechanism for a trans eQTL acting as a master regulator is for it to be a cis eQTL for a transcription factor [41]. In fact, it has been found that a substantial fraction of trans eQTL effects are mediated through a target cis gene [43]. Among the five chromosome 12 SNPs in strong LD that are detected as significant by ADELLE, one was previously shown to be a cis eQTL [4] (see Table S7). Other mechanisms by which a SNP may act on trans genes have been discussed [2] and may be relevant for these SNPs.

## Discussion

For trans-eQTL mapping, in order to meet rigorous standards of genomewide significance, the common strategy of considering the entire set of p-values for testing each SNP against each trans trait requires a severe multiple testing correction, because both SNP-wise and trait-wise correction is required. The resulting threshold is too strict for anything other than extremely strong associations to pass. Since a trans-eQTL association signal is not expected to be particularly large, this strategy does not seem well-suited to detecting trans-eQTLs. A global testing strategy in which association test statistics for a single SNP are combined across multiple expression traits into a single test statistic for each SNP has the potential help alleviate this problem because the resulting global test p-values need only be corrected for the number of SNPs. Whether a global test actually represents an improvement can depend entirely on the form of the global test. For example, the global test based on Min-*p* which is one of the methods considered in our simulations is essentially the same as the common strategy.

We have developed a global testing method ADELLE that is tailored for trans-eQTL mapping. ADELLE is designed to have high power when a trans-eQTL is associated with multiple expression traits, where the proportion of associated traits is small as a subset of all traits tested, and where the individual effect sizes may be relatively weak. We have shown through simulation studies and a reanalysis of a mouse AIL data set that our method, ADELLE, is able to detect significant trans eQTL signal that would otherwise not be detected when only individual SNP-trait p-values are considered.

In our simulations, ADELLE was the only method that consistently maintained high relative power when the number of associated expression traits represented .1%–2% of the total number of traits tested, and it had significantly higher power than the other methods when the number of associated expression traits represented around 0.2%–1% of the total number of traits tested. These are particularly relevant ranges for trans eQTLs because it is expected that they will often be associated with many, rather than just a single, gene. In fact, as seen in our analysis of the AIL, ADELLE is able to reject the global null hypothesis even when none of the individual trait *p*-values for a SNP are particularly small (i.e., they do not meet the significance threshold when correcting only for SNP-wise multiple testing, much less the more stringent standard of correcting for both SNP-wise and trait-wise multiple testing). This shows the ability of ADELLE to effectively combine multiple sub-significant association signals for a given SNP to enable genome-wide significant trans-eQTL detection.

ADELLE needs only summary statistics (consisting of (1) either *Z* scores or else p-values and the signs of the estimated effect sizes and (2) a sample correlation matrix for the traits) to perform its analysis. A distinct advantage of a method that only requires summary statistics is the ease with which they can be shared. This is especially relevant in human data where concerns regarding privacy and the risk of re-identification can make the sharing of original, individual level data problematic. In addition, sharing of summary statistics avoids the duplication of computation and effort that results when the original data must go through the process of quality control, normalization, testing, etc. multiple times. Sharing of the summary statistics is not without burden, however. The storage and sharing of summary statistics can be demanding, particularly in trans-eQTL studies where pairwise combinations of SNPs and genes result in a very large number of tests. In practice, even the complete set of *Z* scores may not be available. An advantage of ADELLE is that it is only based on the *qD* most significant results for each SNP, where *q* is set by the user, so a full set of summary statistics is not required. In addition, ADELLE could in principle be modified to use only the summary statistics for tests that meet a certain pre-specified significance level, rather than using a fixed number of top results for each SNP.

ADELLE is implemented in a freely downloadable software package that will be made available at https://www.stat.uchicago.edu/~mcpeek/software/index.html. Computing the ELL statistic at a SNP is not time consuming. This is especially true when a precompute grid is used for the *l*-values. On a desktop computer, the precompute grid for this study took less than one minute to create. Once this grid is created, on the order of 1,000 SNPs can be analyzed per minute. The primary computational burden results from the Monte Carlo approach to determining the null distribution of our statistic. A more efficient approach to determine statistical significance is an area for future work.

Understanding the underlying biological mechanisms of trans acting effects on gene expression is a challenging task that will involve combining evidence from various lines of investigation. Here we focused on the statistical problem of identifying SNPs that affect variation in gene expression of distant genes. The combination of relatively weak effects with a very large number of tests make this a particularly difficult problem. The statistical methodology we developed for this problem, however, is general and can easily be applied to a larger set of common problems in genomics. Most any problem that involves an aggregating, or a set-based, test may benefit from our approach. For instance, tests of gene sets, SNP sets, and pathways fall into this category as do phenome wide association tests and tests which involve potential interactions when there are many possibly interacting variables, such as epistasis. In fact, as technology in the field of genomics progresses, and the number of variables, conditions and contexts grows with the size of data sets, we expect highly sensitive methods such as ADELLE to be a valuable tool in the process of developing deeper insights from the data.

## Acknowledgments

We gratefully acknowledge N. Gonzales for her help with the AIL data set. This work was funded by NIH grant R01 HG001645 to MSM.

## Notes

### Competing Interest Statement

The authors have declared no competing interest.

### Summary of Updates

Revised the methods, simulations and data analysis and made wording changes throughout.

