## Supplementary text and tables for "ADELLE: A global testing method for Trans-eQTL mapping"

### S1 Text

#### Model for $Z$

In an eQTL mapping study in which $\tilde{D}$ expression traits and $M$ genome-wide SNPs are observed on each of $N$ individuals, define $G$ to be the $M \times N$ genotype matrix for the SNPs and $Y$ to be the $\tilde{D} \times N$ phenotype matrix for the expression traits. Let $Z$ denote the $\tilde{D} \times M$ matrix of test statistics, where $Z_{dm}$, the $(d, m)$th entry of $Z$, is the test statistic for association between trait $d$ and SNP $m$, and where each $Z_{dm}$ is assumed to be standard normal under the null hypothesis of no association between trait $d$ and SNP $m$. A typical situation would be that $N \ll \min(\tilde{D}, M)$, i.e., there are many more expression traits and SNPs than there are individuals in the study.

In the simplest case, suppose that there are neither confounding covariates nor important population structure in the trait models. Then under the null hypothesis that none of the SNPs are eQTLs for any of the traits, the $N$ columns of $Y$ are assumed to be i.i.d. draws from $N_{\tilde{D}}(\mu_{trait}, V_{trait})$, where $\mu_{trait}$ is the $\tilde{D} \times 1$ vector of trait means and $V_{trait}$ is the $\tilde{D} \times \tilde{D}$ trait covariance matrix, and the $N$ columns of $G$ are assumed to be independent, each having mean vector $\mu_{geno}$ of length $M$ and $M \times M$ covariance matrix $V_{geno}$. Furthermore, suppose that $Z_{dm}$ is obtained as the t-statistic for testing significance of SNP $m$ in a simple linear regression of trait $d$ on SNP $m$, for $1 \leq d \leq \tilde{D}, 1 \leq m \leq M$. Then for reasonably large $N$ (even when $N \ll \min(\tilde{D}, M)$), the covariance structure of the elements of $Z$ is given by

$$\text{Cov}(\text{vec}(Z)) = C_{geno} \otimes C_{trait}, \tag{1}$$

where $C_{geno}$ is the correlation matrix derived from $V_{geno}$ and $C_{trait}$ is the correlation matrix derived from $V_{trait}$. A consequence of this model is that each row of Z has covariance matrix $C_{geno}$ and each column of $Z$ has covariance matrix $C_{trait}$. In the more general setting with a linear mixed model for the trait, including covariates, the model in Eq 1 becomes approximate rather than exact.

Since each element of $Z$ is marginally standard normal and they have the correlation structure in Eq 1, it would be tempting to suppose that the joint distribution of the elements of $Z$ must be multivariate normal, and to suppose that therefore $Z$ must have the matrix normal distribution $\text{MN}_{\tilde{D},M}(\mathbf{0}, C_{trait}, C_{geno})$. However, this is false. In fact the distribution of $Z$ is a mixture of matrix normal distributions, where each component of the mixture has mean 0 and variance 1 for every element of $Z$, but where different mixture components have different correlation matrices for vec($Z$). One way to specify the distribution of $Z$ in this case is to note that conditional on $G$, $Z$ does have a matrix normal distribution in large samples:

$$Z|G \sim \mathrm{MN}_{\tilde{D},M}(\mathbf{0}, C_{trait}, \hat{C}_{geno}) \tag{2}$$

where $\hat{C}_{geno}$ is the $M \times M$ sample correlation matrix of the given $G$, which is a matrix of rank $N-1$. From this formulation, it can be seen that each $Z$ is only of rank $N-1$, which makes sense, because there are only $N$ observations of each phenotype and genotype on which the entire matrix $Z$ is based. It can also be seen that the unconditional distribution of $Z$ will be a mixture of matrix normals, where we mix over $G$. We note that it is also true in this case that

$$Z|Y \sim \mathrm{MN}_{\tilde{D},M}(\mathbf{0}, \hat{C}_{trait}, C_{geno}), \tag{3}$$

where $\hat{C}_{trait}$ is the $\tilde{D} \times \tilde{D}$ sample correlation matrix of the given $Y$, so $\hat{C}_{trait}$ is of rank $N-1$.

In this simple setting, $\Omega = C_{trait}$ where $\Omega$ is as defined in the main text (e.g. main text Eq (5)), and we could directly estimate $C_{trait}$ by first forming the sample correlation matrix $\hat{C}_{trait}$ for the $\tilde{D}$ traits. However, because typically $N \ll \tilde{D}$, the estimate $\hat{C}_{trait}$ is low rank. We, therefore, regularize it by using the shrinkage estimator [1] $\widehat{\Omega} = w\hat{C}_{trait} + (1-w)I$ as described below in subsection **Regularization of sample covariance matrix**.

Other approaches are possible. For example, one could consider a more general null model for $Z$:

$$Z|G \sim \mathrm{MN}_{\tilde{D},M}(\mathbf{0}, \Omega, C_G), \tag{4}$$

where $C_G$ is a $M \times M$ correlation matrix that is a function of $G$ and has rank $N-1$ (or, more generally, rank($C_G$) $= N-k-1$ if $k$ PCs, PEER factors, or other covariates have been regressed out of the expression traits), and $\Omega$ is a $\tilde{D} \times \tilde{D}$ correlation matrix, where rank($\Omega$) = rank($C_{trait}$), which we assume is equal to $\tilde{D}$. Properties of this model are that $Z$ is of rank $N-k-1$, every column of $Z$ has covariance matrix $\Omega$ and every row of $Z$ has covariance matrix $C_G$.

A simple (but low-rank) estimator for $\Omega$ in this model would be the sample correlation matrix for the rows of $Z$, given by cov2cor($Z(I_M - M^{-1}\mathbf{1}_M\mathbf{1}_M^T)Z^T$, where cov2cor is the function that maps a symmetric positive semi-definite matrix $A$ with positive diagonal elements to a matrix of the same size with $(i,j)$th element $A_{ij}/\sqrt{A_{ii}A_{jj}}$. If $C_G$ were available, a more efficient (but still low-rank) estimator could be obtained by both decorrelating and centering the rows of $Z$ to obtain cov2cor($Z(C_G^- - C_G^-\mathbf{1}_M(\mathbf{1}_M^T C_G^-\mathbf{1}_M)^{-1}\mathbf{1}_M^T C_G^-)Z^T$), where $C_G^-$ is the Moore-Penrose generalized inverse of the (singular) matrix $C_G$. More generally, suppose we used matrix $\tilde{C}^-$ in the estimator $\hat{\Omega}_1 = \text{cov2cor}(Z\tilde{C}^- Z^T)$, where $\tilde{C}^-$ is a symmetric, positive semi-definite matrix.

To decide which choice of $\tilde{C}^-$ to use in the estimator $\hat{\Omega}_1$, we could consider the following simple setting for estimation of a single correlation value $\rho$. Suppose $(X_1, Y_1), \ldots, (X_n, Y_n)$ are i.i.d. bivariate normal with mean vector $(0, 0)$ and with $2 \times 2$ covariance matrix having both diagonal elements equal to 1 and both off-diagonal elements equal to $\rho$. Then, if we let $\rho_{d1,d2}$ denote the $(d1, d2)$th entry of $\Omega$, where $d1 \neq d2$, in large samples the $(d1, d2)$th entry of $\hat{\Omega}_1$ would be approximately unbiased for $\rho_{d1,d2}$ with variance $(1 - \rho^2_{d1,d2})^2 \text{tr}(C_G \tilde{C}^- C_G \tilde{C}^-)/[\text{tr}(C_G \tilde{C}^-)]^2$, which is minimized when $\tilde{C}^- = C_G^-$, in which case it becomes $(1 - \rho^2_{d1,d2})^2/\text{rank}(C_G)$. However, to increase the robustness of the estimator, we would usually prefer to choose a $\tilde{C}^-$ that is orthogonal to $1_M$, so that we are also centering $Z$. Within this class of estimators, the variance is minimized when $\tilde{C}^- = C_G^- - C_G^- \mathbf{1}_M (\mathbf{1}_M^T C_G^- \mathbf{1}_M)^{-1} \mathbf{1}_M^T C_G^-$, which leads to the second low-rank estimator suggested above. The resulting variance of the estimator is $(1 - \rho^2_{d1,d2})^2/(\text{rank}(C_G) - 1)$.

Practical choices one could consider would be $\tilde{C}^- = I - M^{-1} 1_M 1_M^T$ (which makes $\hat{\Omega}_1$ the sample correlation matrix) or $\tilde{C}^- = \hat{C}^-_{geno} - \hat{C}^-_{geno} \mathbf{1}_M (\mathbf{1}_M^T \hat{C}^-_{geno} \mathbf{1}_M)^{-1} \mathbf{1}_M^T \hat{C}^-_{geno}$, where $\hat{C}_{geno}$ is the sample correlation matrix of $G$, which would be the optimal choice among all $\tilde{C}^-$ that are orthogonal to $1_M$, in the special case of simple linear regression. Given $\hat{\Omega}_1$, which is generally of rank $N - k - 1$, we could apply regularization to obtain full-rank $\widehat{\Omega}$ as in the next subsection. However, a major drawback of this alternative approach to estimating $\widehat{\Omega}$ is that the entire matrix $Z$ is required, while in practice, summary statistics are often only available for a subset of the $Z$ matrix. Because of this drawback, we do not give further consideration to the alternative approach. In contrast, the $\Omega$ estimation approach described in the main text is more widely applicable, because it can be used when only a subset of the $Z$ matrix is available, as long as the available summary statistics include the sample covariance of the trait values.

##### Regularization of sample covariance matrix

To estimate $\Omega$, we first directly estimate $C_{trait}$ by forming the sample correlation matrix $\hat{C}_{trait}$ for the $\tilde{D}$ traits. However, because typically $N \ll \tilde{D}$, the estimate $\hat{C}_{trait}$ will be low rank. We, therefore, regularize it by using the shrinkage estimator [1] $\widehat{\Omega} = w\hat{C}_{trait} + (1 - w)I$. We now describe how we choose the regularization parameter $w$.

For the beta-binominal approximation defined in the next subsection, from Equation 10 it can be seen that the leading term of the beta-binomial approximation to the quantity $V_0[S(c)]$ needed for $l$-value calculation depends on $\widehat{\Omega}$ through the quantity $O_2(\widehat{\Omega})$, where for any $D \times D$ positive semi-definite matrix $M$ we define $O_2(M) = \sum_{1 \leq k < l \leq D} (M_{kl})^2$. Therefore, in our regularization procedure, we focus on choosing $w$ that leads to accurate recovery of $O_2(\hat{C}_{trait})$ in simulated replicates.

We perform a search over $w \in (0, 1)$, and for each choice of $w$, we perform the following steps: (1) form

$\widehat{\Omega} = w\hat{C}_{trait} + (1-w)I_D$; (2) sample 4 replicates of the $D \times N$ matrix $Y$ from $MN(0, \widehat{\Omega}, I_M)$ to obtain $Y^{(1)}, Y^{(2)}, Y^{(3)}$, and $Y^{(4)}$; (3) for $i = 1, 2, 3, 4$, let $\hat{C}^{(i)}$ denote the sample trait covariance matrix based on replicate $Y^{(i)}$; (4) compare $O_2(\hat{C}_{trait})$ to $.25 \sum_{i=1}^{4} O_2(\hat{C}^{(i)})$, and choose $w$ to minimize the absolute difference between these two quantities.

#### Beta-binomial approximation

In this subsection, we describe the beta-binomial approximation [2] used to calculate values of $F_{(d)}$, the cdf of $\pi_{(d)}$, for $d = 1, \ldots, qD$, which is needed for obtaining the l-values used in forming the ADELLE test statistic. From Eq (7) in the main text, we have for the $l$-value

$$l_d(h) \equiv F_{(d)}(h) \equiv P_0(\pi_{(d)} \leq h) = P_0(S(-\Phi^{-1}(h/2)) \geq d), \tag{5}$$

where $S(c) = \sum_{d=1}^{D} \mathcal{I}\{|Z_d| \geq c\}$. If $\Omega = I$, then for $c \geq 0$, $S(c)$ has the null distribution of a Binomial$(D, 2\Phi(-c))$ random variable, and

$$F_{(d)}(h) = 1 - \sum_{j=0}^{d-1} \binom{D}{j} h^j (1-h)^{D-j}, \quad h \in [0,1], \tag{6}$$

which is the CDF of a Beta$(d, D+1-d)$ distribution. When $\Omega \neq I$, the null mean of $S(c)$ will remain the same, $E_0[S(c)] = 2D\Phi(-c)$, however the null variance is

$$\begin{aligned} V_0(S(c)) &= V_0\left(\sum_k \mathcal{I}\{|Z_k| \geq c\}\right) \\ &= 2D\Phi(c)(1-2\Phi(c)) + 2\sum_{i=1}^{D-1}\sum_{j=i+1}^{D} \operatorname{cov}\left(\mathcal{I}\{|Z_i| \geq c\}, \ \mathcal{I}\{|Z_j| \geq c\}\right) \\ &> 2D\Phi(c)(1-2\Phi(c)) \end{aligned}$$

Where the last inequality follows from the positive correlation of the magnitudes of any correlated, mean-zero, bivariate Gaussians.

Following the method proposed by [2] (see also [3]), we approximate the distribution of $S(c)$ with a beta-binomial distribution $BB(D, a, b)$. The beta-binomial is the distribution formed from the binomial distribution when the success probability is drawn from a beta distribution. It is convenient to reparameterize the beta-binomial in terms of parameters $\lambda = \frac{a}{a+b}$ and $\gamma = \frac{1}{a+b}$. If $X$ is drawn from a BB($D$, $\lambda$, $\gamma$), where

$\lambda > 0$, $\gamma > 0$, and $D$ is a positive integer, then $X$ has the following properties:

$$f_{D,\lambda,\gamma}(x) \equiv P(X = x) = \binom{D}{x} \frac{B(\frac{\lambda}{\gamma} + x, \frac{1-\lambda}{\gamma} + D - x)}{B(\frac{\lambda}{\gamma}, \frac{1-\lambda}{\gamma})}, \tag{7}$$

$$\lambda = E(X)/D, \text{ and} \tag{8}$$

$$\frac{\gamma}{1+\gamma} = \frac{V(X) - D\lambda(1-\lambda)}{D(D-1)\lambda(1-\lambda)}, \tag{9}$$

where $B(a, b)$ is the beta function.

To approximate the distribution of $S(c)$ we choose $\lambda$ and $\gamma$ by the method of moments. Barnett et al. [2] derived the following:

$$\begin{aligned} E_0\left[S\left(c\right)\right] &= 2D\Phi(-c) \\ V_0\left[S\left(c\right)\right] &= D\left[2\overline{\Phi}(c) - 4\overline{\Phi}^2(c)\right] + 4D(D-1)\phi^2(c)\textstyle\sum_{i=1}^{\infty} \mathcal{H}_{2i-1}^2(c)\rho(2i)/(2i)! \end{aligned} \tag{10}$$

Where $\mathcal{H}_i$ are the Hermite polynomials, $\rho(i) = \frac{2}{D(D-1)} \sum_{1 \leq k < l \leq D} \left(\Omega_{kl}\right)^i$, where $\Omega_{kl}$ is the $(k, l)$th entry of $\Omega$, and $\phi$ and $\overline{\Phi}$ are the density and survivor functions of a standard normal, respectively. Note that Eqs 8, 9, and 10 imply that once $\lambda$ is set by the method of moments, knowing $\lambda$ determines $c$ which determines $V_0\left[S\left(c\right)\right]$ and thus determines $\gamma$. So we will drop the writing of $\gamma$ (and $D$) below to simplify notation, and define $f_\lambda(x)$ to be the beta-binomial density with parameters $(D, \lambda, \gamma)$, where $\lambda$ and $\gamma$ are related by the equations $\lambda = 2\Phi(-c)$ and

$$\frac{\gamma}{1+\gamma} = \frac{V_0[S(c)] - D\lambda(1-\lambda)}{D(D-1)\lambda(1-\lambda)},$$

with $V_0[S(c)]$ given in 10.

Applying this to Eq 5,

$$
\begin{aligned}
l_d(h) \equiv F_{(d)}(h) &= P_0\left[\pi_{(d)} \le h\right] \\
&= P_0\left[S\left(-\Phi^{-1}\left(h/2\right)\right) \ge d\right] \\
&= 1 - \sum_{k=0}^{d-1} P_0\left[S\left(-\Phi^{-1}\left(h/2\right)\right) = k\right] \\
&\approx 1 - \sum_{k=0}^{d-1} f_h\left(k\right) \\
&= \sum_{k=d}^{D} \binom{D}{k} \frac{B(\frac{\lambda}{\gamma}+k, \frac{1-\lambda}{\gamma}+D-k)}{B(\frac{\lambda}{\gamma}, \frac{1-\lambda}{\gamma})} \\
&\equiv \hat{F}_{(d)}(h) \equiv \hat{l}_d(h). \qquad (11)
\end{aligned}
$$

#### Pre-computation for the ADELLE test

Our ADELLE method lends itself to a pre-computation to reduce computation time when it will be applied to a large number of SNPs and a large number of traits. Suppose we observe $M$ SNPs along with the $D$ traits. Then, from Equation (11), calculating $M$ ADELLE statistics involves $M \times \sum_{j=1}^{qD}(D-j+1) \approx MqD^2$ evaluations of a beta-binomial probability mass function $f_h(k)$. However, we also see that computation of $\hat{l}_1(h)$ necessarily also computes all the $l$-values $\hat{l}_1(h), \ldots, \hat{l}_D(h)$.

This motivates a computationally efficient strategy for approximating the function $\hat{F}_{(d)}(h)$ for arbitrary $h$ and $1 \le d \le qD$ by pre-computing a $qD \times H$ matrix of $l$-values, where $H$ is the number of values of $h$ in the grid. We choose a set of pre-computation points, $\mathcal{H} = \{h_1, \ldots, h_H\}$, where $0 < h_1 < \cdots < h_H = 1$, and evaluate each $\hat{l}_d$ only on the points in $\mathcal{H}$. For $1 \le d \le qD$ but $h \notin \mathcal{H}$ we linearly interpolate when $h_1 < h < h_H$,

$$
\hat{l}_d(h) \approx \hat{l}_d(h_i) + \frac{\hat{l}_d(h_{i+1}) - \hat{l}_d(h_i)}{h_{i+1} - h_i}(h - h_i)
$$

where $h_i < h < h_{i+1}$. For $h < h_1$ we compute the $l$-value directly from (11).

In general both the computation time and the accuracy of the approximation tend to increase with the number of points $H$ and the density of the pre-compute grid. In the data analysis, we use $H = 10^5$ and $h_1 = 10^{-20}$. A naive means of assigning the values of $\mathcal{H}$ would be to create an evenly spaced grid between $h_1$ and 1, but this devotes few points, and little resolution to small values of $h$. Instead we choose $\mathcal{H}$ to be the geometric sequence $h_i = h_1^{\frac{H-i}{H-1}}$ for $1 \le i \le H$. This results in 5,000 grid points for each order of magnitude, scaled logarithmically, for our choice of $h_1$ and $H$.

##### Generation of the correlation matrix for the simulations

We generated a random correlation matrix to serve as the $D \times D$ correlation matrix between traits. We created this matrix by generating a random orthogonal matrix to serve as eigenvectors and sampling a set of eigenvalues from a gamma distribution with shape and scale parameters of 1.84 and 0.037, respectively. A value of 0.5 was added to all the eigenvalues to prevent any values being close to zero. The eigenvectors and eigenvalues are combined to form a covariance matrix which we then convert to a correlation matrix. These parameters of the gamma distribution were chosen such that when a sample of size 208 was taken, the distribution of off diagonal values was similar to that seen in the AIL expression data correlation matrix.

#### Supporting Tables

**Table S1. Power comparison for trans eQTL mapping when number of associated traits = 5**

| | Power (s.e.) at significance level | | | |
|---|---|---|---|---|
| Method | 0.05 | 0.01 | 0.001 | 1e-04 |
| Adelle | 0.717 (0.014) | 0.501 (0.016) | 0.281 (0.014) | 0.113 (0.010) |
| Cauchy | *0.813 (0.012) | *0.643 (0.015) | *0.353 (0.015) | *0.172 (0.012) |
| Simes | *0.794 (0.013) | *0.618 (0.015) | *0.339 (0.015) | *0.167 (0.012) |
| min-P | *0.790 (0.013) | *0.612 (0.015) | *0.336 (0.015) | *0.166 (0.012) |
| G-null | 0.608 (0.015) | 0.344 (0.015) | 0.131 (0.011) | 0.035 (0.006) |
| Sum-$\chi^2$ | 0.122 (0.010) | 0.035 (0.006) | 0.006 (0.002) | 0.000 (0.000) |
| CPMA | 0.062 (0.008) | 0.020 (0.004) | 0.002 (0.001) | 0.000 (0.000) |

Power is based on 1000 simulated replicates. s.e. denotes standard error of the estimate. A starred number denotes the highest power attained or power that is not significantly different (based on a z-test at level .05) from the highest power attained by any of the methods.

**Table S2. Power comparison for trans eQTL mapping when number of associated traits = 10**

| | Power (s.e.) at significance level | | | |
|---|---|---|---|---|
| Method | 0.05 | 0.01 | 0.001 | 1e-04 |
| Adelle | *0.617 (0.015) | *0.369 (0.015) | *0.182 (0.012) | *0.063 (0.008) |
| Cauchy | *0.657 (0.015) | *0.405 (0.016) | *0.169 (0.012) | *0.055 (0.007) |
| Simes | *0.632 (0.015) | *0.388 (0.015) | *0.162 (0.012) | *0.055 (0.007) |
| min-P | *0.614 (0.015) | *0.381 (0.015) | *0.160 (0.012) | *0.055 (0.007) |
| G-null | 0.517 (0.016) | 0.233 (0.013) | 0.080 (0.009) | 0.016 (0.004) |
| Sum-$\chi^2$ | 0.148 (0.011) | 0.042 (0.006) | 0.008 (0.003) | 0.000 (0.000) |
| CPMA | 0.082 (0.009) | 0.025 (0.005) | 0.004 (0.002) | 0.000 (0.000) |

Power is based on 1000 simulated replicates. s.e. denotes standard error of the estimate. A starred number denotes the highest power attained or power that is not significantly different (based on a z-test at level .05) from the highest power attained by any of the methods.

**Table S3. Power comparison for trans eQTL mapping when number of associated traits = 20**

| Method | Power (s.e.) at significance level 0.05 | 0.01 | 0.001 | 1e-04 |
|---|---|---|---|---|
| Adelle | *0.724 (0.014) | *0.488 (0.016) | *0.203 (0.013) | *0.063 (0.008) |
| Cauchy | 0.675 (0.015) | 0.371 (0.015) | 0.127 (0.011) | 0.040 (0.006) |
| Simes | 0.617 (0.015) | 0.343 (0.015) | 0.122 (0.010) | 0.040 (0.006) |
| min-P | 0.597 (0.016) | 0.336 (0.015) | 0.120 (0.010) | 0.039 (0.006) |
| G-null | 0.641 (0.015) | 0.296 (0.014) | 0.082 (0.009) | 0.014 (0.004) |
| Sum-$\chi^2$ | 0.212 (0.013) | 0.069 (0.008) | 0.018 (0.004) | 0.000 (0.000) |
| CPMA | 0.119 (0.010) | 0.036 (0.006) | 0.007 (0.003) | 0.000 (0.000) |

Power is based on 1000 simulated replicates. s.e. denotes standard error of the estimate. A starred number denotes the highest power attained or power that is not significantly different (based on a z-test at level .05) from the highest power attained by any of the methods.

**Table S4. Power comparison for trans eQTL mapping when number of associated traits = 50**

| Method | Power (s.e.) at significance level 0.05 | 0.01 | 0.001 | 1e-04 |
|---|---|---|---|---|
| Adelle | *0.737 (0.014) | *0.472 (0.016) | *0.204 (0.013) | *0.047 (0.007) |
| Cauchy | 0.519 (0.016) | 0.230 (0.013) | 0.040 (0.006) | 0.009 (0.003) |
| Simes | 0.461 (0.016) | 0.196 (0.013) | 0.038 (0.006) | 0.009 (0.003) |
| min-P | 0.433 (0.016) | 0.185 (0.012) | 0.038 (0.006) | 0.009 (0.003) |
| G-null | 0.664 (0.015) | 0.347 (0.015) | 0.087 (0.009) | 0.013 (0.004) |
| Sum-$\chi^2$ | 0.365 (0.015) | 0.155 (0.011) | 0.042 (0.006) | 0.008 (0.003) |
| CPMA | 0.221 (0.013) | 0.084 (0.009) | 0.025 (0.005) | 0.004 (0.002) |

Power is based on 1000 simulated replicates. s.e. denotes standard error of the estimate. A starred number denotes the highest power attained or power that is not significantly different (based on a z-test at level .05) from the highest power attained by any of the methods.

**Table S5. Power comparison for trans eQTL mapping when number of associated traits = 100**

| Method | Power (s.e.) at significance level 0.05 | 0.01 | 0.001 | 1e-04 |
|---|---|---|---|---|
| Adelle | *0.914 (0.009) | *0.713 (0.014) | *0.394 (0.015) | *0.137 (0.011) |
| Cauchy | 0.570 (0.016) | 0.223 (0.013) | 0.051 (0.007) | 0.011 (0.003) |
| Simes | 0.482 (0.016) | 0.196 (0.013) | 0.049 (0.007) | 0.011 (0.003) |
| min-P | 0.443 (0.016) | 0.191 (0.012) | 0.048 (0.007) | 0.011 (0.003) |
| G-null | 0.870 (0.011) | 0.609 (0.015) | 0.262 (0.014) | 0.063 (0.008) |
| Sum-$\chi^2$ | 0.644 (0.015) | 0.381 (0.015) | 0.151 (0.011) | 0.048 (0.007) |
| CPMA | 0.469 (0.016) | 0.236 (0.013) | 0.081 (0.009) | 0.029 (0.005) |

Power is based on 1000 simulated replicates. s.e. denotes standard error of the estimate. A starred number denotes the highest power attained or power that is not significantly different (based on a z-test at level .05) from the highest power attained by any of the methods.

**Table S6. Power comparison for trans eQTL mapping when number of associated traits = 200**

| Method | Power (s.e.) at significance level 0.05 | 0.01 | 0.001 | 1e-04 |
|---|---|---|---|---|
| Adelle | *0.691 (0.015) | *0.392 (0.015) | *0.142 (0.011) | *0.034 (0.006) |
| Cauchy | 0.246 (0.014) | 0.070 (0.008) | 0.011 (0.003) | 0.002 (0.001) |
| Simes | 0.213 (0.013) | 0.065 (0.008) | 0.011 (0.003) | 0.002 (0.001) |
| min-P | 0.198 (0.013) | 0.061 (0.008) | 0.011 (0.003) | 0.002 (0.001) |
| G-null | *0.680 (0.015) | *0.381 (0.015) | *0.136 (0.011) | 0.024 (0.005) |
| Sum-$\chi^2$ | 0.633 (0.015) | *0.369 (0.015) | *0.154 (0.011) | *0.044 (0.006) |
| CPMA | 0.487 (0.016) | 0.266 (0.014) | 0.088 (0.009) | *0.035 (0.006) |

Power is based on 1000 simulated replicates. s.e. denotes standard error of the estimate. A starred number denotes the highest power attained or power that is not significantly different (based on a z-test at level .05) from the highest power attained by any of the methods.

**Table S7. Significant trans eQTL detections by ADELLE in a region of Chrom 12**

| rsID | Chrom | BP | Adelle pvalue | min-p | SNP annotation | cis eQTL | cis gene |
|---|---|---|---|---|---|---|---|
| rs262318378 | 12 | 72948661 | 1.0e-07 | 0.000372 | downstream gene variant | No | NA |
| rs29176728 | 12 | 73012360 | 2.2e-06 | 0.006130 | NA | No | NA |
| rs29223131 | 12 | 73320679 | 2.2e-06 | 0.001360 | intron variant | No | NA |
| rs29197865 | 12 | 73409987 | 1.9e-06 | 0.002200 | upstream gene variant | Yes | Ppm1a |
| rs213877154 | 12 | 73527315 | 2.2e-06 | 0.004000 | NA | No | NA |

min-p is the p-value Bonferroni corrected for the number of traits tested; SNP annotations are from [4]; cis eQTL is whether the SNP is also a cis eQTL; cis gene is the gene target of the cis eQTL.
